# Practices and trends in last-mile delivery of poultry vaccines in rural areas in developing countries: the case of Newcastle disease vaccine delivery in Bungoma County, Kenya

**DOI:** 10.1101/2021.02.19.431949

**Authors:** JK. Chemuliti, KO. Ogolla, SG. Mbogoh, KM. Mochabo, BK Kibore

## Abstract

Newcastle disease (ND) is the single most important infection of village chicken in smallholder farming systems in developing countries. Vaccines for ND control are available but the delivery of safe and potent vaccines in resource-poor settings remains a big challenge due to difficulties in the maintenance of cold chain. This paper reports the results of a study that was carried out in Kenya to assess the storage and handling practices of Newcastle disease vaccines by agro-veterinary shops (agro-shops) during acquisition, storage, and sale to smallholders’ farmers. Data were collected from one hundred and thirty-two agro-shops using semi-structured questionnaires, observation sheets and actual purchase of vaccines over the counter. The results showed that the majority (82 percent) of the agro-shops had a domestic refrigerator that was connected to the electricity grid but many (61 percent) did not have power backup. Sixty percent of them only stocked thermolabile vaccines. Recurrent power outages (62 percent), high cost of electricity (62 percent), and long-distance to vaccine sources (33 percent) were the most common challenges in vaccine storage and sale. Some agro-shops switched refrigerators on and off while others removed vaccines from refrigerators for overnight stay in cool boxes to minimize electricity costs. In some cases, the sale of vaccines was restricted to market days and late afternoon when ambient temperatures were lower to minimize vaccines storage time and vaccine spoilage respectively. Thermostable vaccines were not stored as recommended by the manufacturer and few agro-shops (23 percent) sold reconstituted vaccines. Most shops adequately packaged thermolabile vaccines in improvised materials during sale. Overall, most of the ND vaccine handling and storage practices in the last mile appeared to aim at safeguarding the safety and potency of vaccines, but further research could elucidate the effects of these practices on the quality and potency of ND vaccines.

## Introduction

Newcastle disease (ND) is the single most important infection of village chickens in smallholder farming systems in developing countries (1,2). The disease causes high morbidity and mortality in chickens, thus denying farmers access to income and a cheap and readily available source of animal protein that would contribute to improvements in livelihoods. Vaccines for the control of ND have been developed and are widely available (3,4). It has been empirically demonstrated that these vaccines are effective in reducing morbidity and mortality in chickens and can result in increases in flock sizes and egg production with significant benefits of enhanced income and nutrition to smallholder farmers (2,5–9). However, despite these benefits, the use of poultry vaccines in indigenous poultry is not widespread and remains low in many parts of sub-Saharan Africa. The constraints that hinder the adoption of ND vaccines among smallholder chicken producers have been well enumerated in the empirical literature. Among these, vaccine inaccessibility remains existential due to the lack of cost-effective and efficient vaccine delivery systems in many rural areas (7,10,11). With the development and availability of avirulent thermostable ND vaccines that are less sensitive to cold chains, it was anticipated that vaccine accessibility and utilization among smallholder farmers in rural areas would increase. Initial studies conducted in some parts of Africa showed that the vaccine was effective in controlling the ND, thus leading to improvements in chicken production, food security, and women’s economic status (12,13). However, a recent study in Tanzania showed that the thermostability of ND vaccines was not a major consideration that influenced the purchase decision of the vaccines among smallholder farmers (9).

Several interventions aimed at enhancing the accessibility of poultry vaccines among smallholder farmers have been introduced in many parts of sub-Saharan Africa (14–16). A major focus of such initiatives has been on the last-mile delivery of vaccines leveraging community-based vaccinators to drive improvements in vaccine coverage. Relatively less attention has been directed at the integrity of vaccines at points of sale, commonly the agro-veterinary shops (hereinafter referred to as agro-shops) that stock and sell vaccines to farmers. These agro-shops some of which are located in small towns in rural areas serve as the end of the cold chain where vaccines are stored before they are sold to farmers. While efforts to increase the accessibility of vaccines to smallholder farmers are important, equally important but often overlooked is the storage and handling of the vaccines in these shops which influences their quality and potency. Agro-shops are a critical node in the vaccine supply chain. Vaccines that reach farmers must be potent and able to offer protection to chickens. Understanding how these agro-shops handle vaccines during acquisition, storage and sale to farmers is therefore important in informing strategies and interventions that are aimed at enhancing the integrity of vaccines thereby contributing to effective control of poultry diseases.

Veterinary vaccines have to be handled with care from production through distribution and storage until they are applied to target animals (17). This is essential in order to obtain optimal potency of vaccine and maximal results from a vaccination. The maintenance of the cold chain during acquisition, transport and storage by the vaccine handlers has been shown to be critically important (18,19). One of the major challenges in the distribution of vaccines in rural areas in developing countries is the limited coverage and reliability of electricity and lack of functional cold chain systems. Ensuring vaccine storage under cold temperatures in these resource-poor settings is not easy because cold chain equipment is often unreliable due to equipment failures, power outages and an unreliable electricity grid (20,21). Moreover, preventive maintenance to avoid equipment failures is rarely executed. Spare parts are often not available and where they are, repair of cold chain equipment can take several months (22–24). As a result, vaccines are often exposed to either heat or freezing, making part of them unusable and resulting in vaccine wastage or impaired vaccine efficacy (25–27).

The quality of vaccines is one of the important factors in the successful control of poultry diseases which in turn depends on the proper storage and handling of vaccines. If a vaccine is stored outside the recommended temperature for a considerable time, its potency will be adversely affected thereby reducing protection from vaccine-preventable diseases. Available evidence shows that farmers prefer and prioritize ND vaccination programs that have a high capacity to protect birds from mortality (28) and this invariably depends on the quality of the vaccine.

This paper reports the results of a study that was carried out to assess the vaccine handling practices during the acquisition, storage and sale of Newcastle disease vaccines by agro-veterinary shops in Bungoma County of Kenya. This Newcastle vaccine delivery study was part of a larger study whose objective was to determine how the productivity and market access of indigenous poultry producers in Bungoma County can be improved. This paper thus adds to the growing body of knowledge on vaccine delivery and use among smallholder farmers in developing countries. It is anticipated that the information provided by this paper will inform efforts aimed at improving the delivery of quality poultry vaccines to smallholder farmers.

## Materials and methods

### Description of the study site

Bungoma County is located on the western side of Kenya and lies between latitude 00 28’ and latitude 10 30’ North of the Equator, and longitude 340 20’ East and 350 15’ East. It covers an estimated area of about 3,032 km^2^. The altitude ranges from 1200 to 4321 meters above sea level. The annual rainfall is approximately between 400 mm and 1800 mm and occurs in two seasons: long rains - March to July and short rains - August to October. The annual temperatures range from 0°C to 32°C. The county is divided into nine sub-counties with a total human population of 1,670,570 persons which roughly translates to a population density of about 552.5 persons per sq. km. (29). The total number of households in the County is about 358,796 with an average household size of 4.6 members. Indigenous chicken production is an important source of livelihood for the majority of households in the County, chicken being the second most important livestock species after cattle and with an estimated value of 596 million. The majority of the chickens are of the indigenous ecotypes and are kept under a free-range system with minimal inputs. The smallholder indigenous chicken production has been a major focus for livelihood improvement through various development initiatives in the county. Kitale town which is located in the neighboring Trans Nzoia County was included in the survey because a preceding baseline survey had shown it was a major source of poultry vaccines in Bungoma County.

### Data collection

Data were collected from agro-shops in Bungoma county and Kitale town which is located in the neighboring Trans Nzoia county. All towns and the major trading and market centers in Bungoma County were visited and those that stocked poultry vaccines were delineated and sampled. Data was collected from the agro-shops using a semi-structured questionnaire that was administered to the shop owner or shop attendant after informed consent was sought and obtained. The questionnaire was designed to capture the socio-economic characteristics of the shop owner, cold chain equipment, vaccine sources and brands, and challenges of cold storage. An observation sheet was used to collect data on vaccine packaging and handling practices through making an actual purchase by using enumerators who posed as smallholder farmers. For each vaccine that was procured, the cost, brand, Newcastle disease strain, expiry date and adequacy of packaging materials were noted and recorded. In this study, a vaccine vial was regarded as adequately packed if it was wrapped together with a sufficient block of ice in non-absorbent packaging material and protected from direct sunlight. Directions on vaccine use were also sought and recorded. All shops were georeferenced. This study was internally approved by the Kenya Agricultural and Research Organization (Ref No. KALRO/1/071).

### Data management and analysis

The collected data was entered in a Microsoft© Excel 2016 spreadsheet. The data was then cleaned by checking for missing and incorrect entries, coded and exported to Statistical Package for Social Sciences© (SPSS version 21) software for analysis. Frequencies, descriptive statistics, cross-tabulations and association tests were then carried out. The level of significance for statistical analysis was set at a p-value equal to or less than 0.05.

## Results

### Profile of agro-veterinary shops

One hundred and thirty-two (132) agro-shops were visited and mapped. The majority (74 percent), were located in Bungoma County. Of these, less than a half (39 percent) stocked ND vaccines. The questionnaire response rate among the vaccine stockists was 82 percent. The majority (67 percent) of the agro-shops had been in operation for more than 5 years, while 22 percent of them between 3 to 5 years and the remaining 12 percent in less than 3 years. Among the vaccine sellers, 25 percent were wholesalers of poultry vaccines to smaller agro-shops. Fig 1 shows the distribution of the agro-veterinary shops in the study sites.

**Fig 1.**
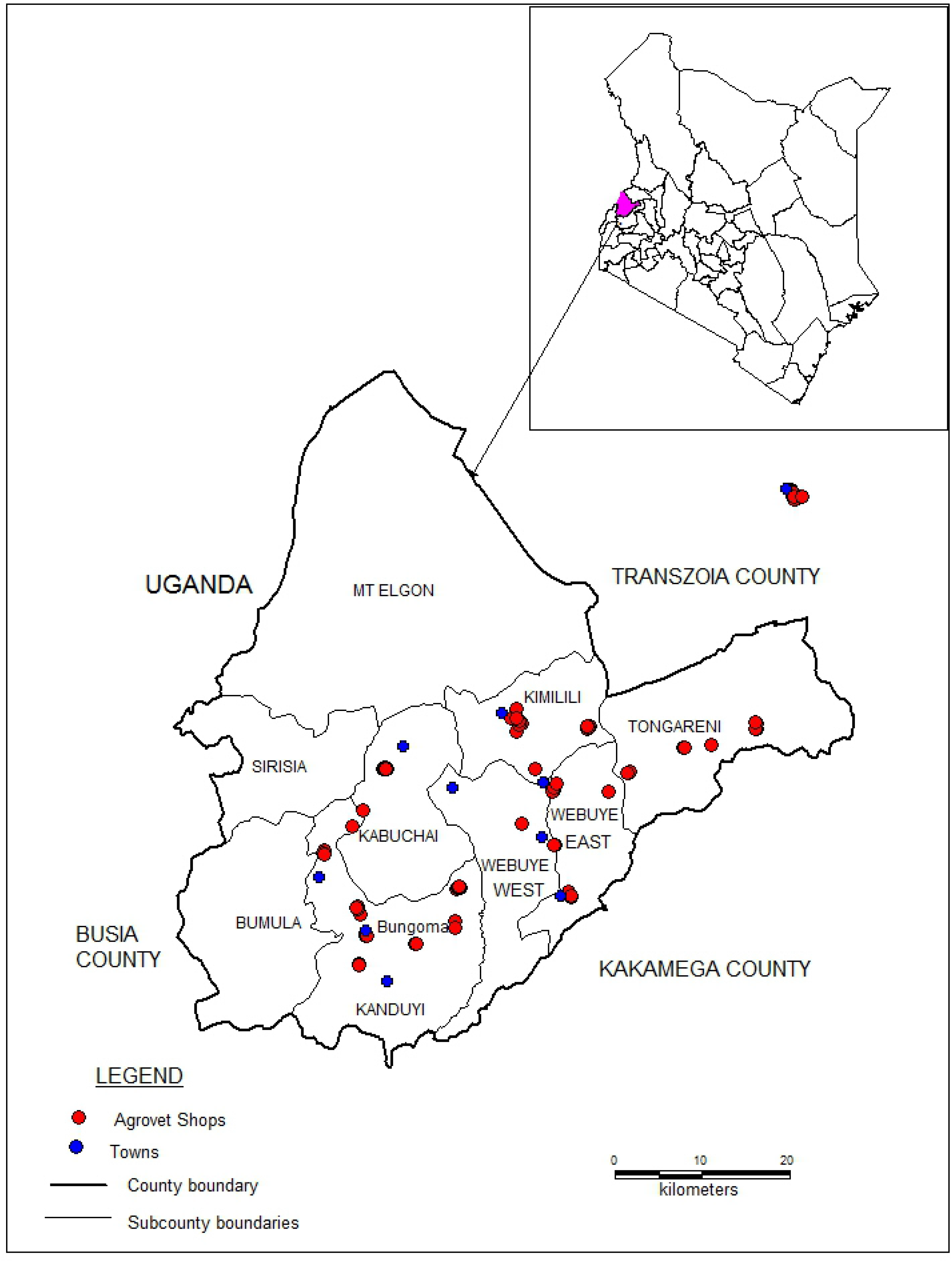
Map of Bungoma showing sampled agro-veterinary shops, towns and trading Centres.

The number and proportion of vaccine stockists were directly related to the proximity to Bungoma town which is the largest poultry market and administrative Centre of Bungoma County. Many (76.5 percent) of the respondents had training in an animal health-related course which is a regulatory requirement of all agro-shops operating in Kenya.

### Characteristics and sources of Newcastle disease vaccines

The agro-shops stocked both thermotolerant and thermolabile Newcastle disease vaccines from different local and international manufacturers. These vaccines included the Lentogenic Lasota strain, F strain, F58 strain and I-2 in freeze-dried tablet and liquid forms. In Kenya, poultry vaccines are mostly supplied by importers and local manufacturers who are located in and around Nairobi, the capital city of Kenya. The vaccines are then distributed to wholesalers and retailers in different parts of the country from where they are sold to a variety of clientele that includes small agro-shops, farmers and animal health service providers. The majority (60 percent) of the agro-shops only stocked thermolabile vaccines, while 15 percent stocked both thermostable and thermolabile types and the remaining 25 percent stocked only the thermostable types. The 100-dose thermolabile ND vaccine vial was the most (81 percent) common dose package and the 25-dose thermostable ND vaccine in a plastic dropper the least (14 percent) common package. The main customers for the ND vaccines were, in order of decreasing frequency, smallholder farmers (92.3%), veterinarians/animal health assistants (50%), commercial chicken farmers (48.1%) and community vaccinators (21.6%).

Fifty-two (52) samples of ND vaccines were procured by enumerators who posed as chicken farmers. These comprised thermolabile freeze-dried forms in small glass vials (69 percent), thermostable liquid forms in plastic droppers (10 percent) and reconstituted vaccines (21 percent). The labels on the procured vaccine vials showed that they were manufactured in India (71 percent), Spain (17 percent) and locally in Kenya (12 percent). All vaccine vials in plastic droppers were manufactured in Kenya. The product labels on all the procured vaccines showed that they required storage temperatures of between +2°C and + 8°C. The average cost of a 100-dose vial was KES 259 (range 220-320). Two of the vaccine vials were found to be beyond their expiry dates.

### Cold chain infrastructure and vaccine storage practices

The majority (94 percent) of the sampled agro-shops that stocked vaccines had cold chain equipment that was connected to the electricity grid. Eighty-two percent of the vaccine stockists had a domestic refrigerator with a freezer compartment in single or double door styles for vaccine storage, 10 percent had both a refrigerator and a standalone freezer, 6 percent had only a standalone freezer and another 6 percent had a cool box where vaccines were stored. Many (61 percent) of the agro-shops did not have a power backup. Power backup consisted of manually operated standby generators (65 percent), solar power system (35 percent) and both the generator and solar system in a few (5 percent) agro-shops. The vaccines were mostly (94 percent) transported from wholesalers to the agro-shops in cool boxes or improvised vaccine packaging materials filled with ice packs to maintain low temperatures. The vaccines were then unpacked and stored in a refrigerator until they were sold. The thermostable vaccines in the plastic droppers were stored in cool boxes without ice packs or on shelves in the agro-shops. The most common challenge that the agro-shops encountered in their endeavor to safely handle and store the vaccines as per the best vaccine storage practices were recurrent power outages (62 percent), high cost of electricity (62 percent) and long-distance to vaccine sources (33 percent). Other constraints that were also mentioned were the high acquisition costs of standby generators and solar power systems, high cost of fuel and maintenance costs for standby generators, and lack of materials for packaging vaccines to customers. Some agro-shops had devised different ways to circumvent the frequent power interruptions by installing standby generators or power solar systems. However, such measures were observed more frequently in more established agro-shops (23 percent) that had been in operation for more than 5 years, mostly located in the towns and larger trading centers in the County. Only a small fraction (11 percent) of agro-shops in more remote areas had power backup. Some agro-shops located in remote areas devised less costly practices, such as switching on and off the refrigerators to minimize power costs. These agro-shops indicated that they switched off the refrigerator at the close of business each day, transferred vaccines into ice-packed cool boxes for an overnight stay and returned them into the refrigerators the next morning. Other agro-shops restricted the sale of vaccines to specific days, usually market days, to minimize vaccine storage time and therefore the cost of having the refrigerator on for a long time. In such cases, vaccines were acquired a day before the market day, kept overnight and the entire stock sold off to customers the next day. In some instances, the sale of vaccines was restricted to specific hours, usually late in the afternoon and mostly after four o’clock when ambient temperatures were lower and only to customers who had thermos flasks to carry the vaccines. This practice was common in agro-shops located in the more remote areas away from major shopping centers.

### Vaccine handling and packaging practices during the sale

Most (61 percent) of the agro-shops sold the vaccines in original vaccine vials as supplied by the manufacturers. A few shops (23 percent), especially those located in more remote areas, sold reconstituted vaccines. A common practice in the sale of thermolabile ND vaccines was for the customer to first specify the number of birds and their locality. The information was then used to determine the number of vaccine doses and adequacy of the packaging materials, in particular the quantity of ice. Vaccine brand, virus strain and expiry date on the vaccine label were not major considerations in the purchase of the vaccines. It was observed that the small size of vaccine vials made it rather difficult for customers to read the information provided on the label. The vaccines were commonly packaged in improvised packaging materials that included, for example, used small plastic containers, polythene bags and small tins. The most common practice by agro-shops was to place the vaccine vial in a used non-absorbent packaging material mostly polythene paper, with a small block of ice which was then secured with cello tape and then wrapped in an opaque piece of paper. Most such shops provided a 10 ml plastic bottle of a diluent, a needle and a syringe alongside the vaccine for a 100-dose vial. In a few cases, vaccine vials were sold in a thermos flask that was filled with a few blocks of ice. An assessment of the adequacy of vaccine packaging revealed that the majority (88 percent) of the agro-shops wrapped ND vaccine adequately for customers. The general advice given to the customers during the sale was to use the vaccine before the ice melts. The agro-shops that sold reconstituted vaccines to their customers dispensed them in syringes or small polythene bags which were wrapped in ice. The volume of the reconstituted vaccine that was dispensed to a customer was dependent on the number of chickens that the customer had indicated. Most of the agro-shops that sold reconstituted vaccines were located in more remote areas in the county. In some areas, vaccines were only sold to customers late in the afternoon, mostly after 4 o’clock when environmental temperatures were lower. Thermostable vaccines in plastic tubing were wrapped in paper and given to the customers without an ice pack.

## Discussion

This study has highlighted vaccine handling practices that were used by agro-shops during the acquisition, storage and sale of Newcastle disease vaccines in the last mile delivery. The agro-shops stocked vaccines that required cold chain temperatures to maintain potency. An assessment of the cold chain equipment in the agro-shops revealed that most of them (82 percent) had domestic refrigerators powered by an electricity grid. However, the existence of agro-shops with cool boxes and freezers only for vaccine storage casts doubt on the quality and potency of the vaccines that they sell to their customers as all the procured vaccines in the study area were indicated for storage of between +2°C and + 8°C. In many low-income countries, vaccine delivery systems have remained largely unchanged due to challenging contextual factors that have limited their ability to meet vaccination program requirements (22). Faced with the persistent challenge of frequent power outages and the rising cost of power, some of the agro-shops in the study area used different strategies to maintain cold storage conditions. Restricting the sale of vaccines to specific days and hours of sale could be regarded as good practice under these circumstances where the maintenance of the cold chain is difficult. However other practices, such as the transferring of vaccines in and out of refrigerators and switching refrigerators on and off to reduces power costs could inadvertently be exposing vaccines to heat and cold and may affect their quality and potency. For a vaccine to be useful in conferring protection against infections, it must be pure, safe, potent, and effective (19). This invariably depends on good vaccine handling practices. This study has revealed that there were few agro-shops (18 percent) that did not observe the basic requirements for safe storage and handling of vaccines. These shops did not have cold chain equipment and kept the vaccines outside the recommended cold chain temperatures. This practice compromises the quality and potency of ND vaccines and undermines the objectives of vaccination. The fact that such agro-shops were located in the deeper rural areas may suggest that they could be operating away from the radar of regulators who are by law required to ensure that all veterinary products are stored under the right conditions for quality and safety.

The risk of vaccine quality deterioration is perhaps greatest in the last mile delivery as vaccines move from agro-shops to farms where they are administered to animals. Vaccines are extremely vulnerable to spoilage when they leave refrigerators as they become more exposed to high temperatures and light. Therefore, the way the vaccine vials are packaged for customers at the point of sale is critical. It is generally recommended that poultry vaccines are carried in portable insulated carriers (cool boxes, vaccine carriers and coolers) with frozen ice packs or locally available packaging materials that can maintain cold chain vaccine temperatures (30 as they move from refrigerator storage to farms for vaccination. An assessment of the adequacy of vaccine packaging materials by this study showed that most agro-shops (88 percent) adequately packaged the ND vaccine with sufficient quantities of ice in a non-absorbent material which was wrapped in an opaque piece of paper to protect it from light. The practice of asking customers where they came from by some agro-shops before packaging the vaccines to determine the adequacy of packing materials and in some cases limiting the sale of vaccines only to customers with thermos flasks are generally good practices that can help maintain vaccine quality and potency.

The recommended cold chain temperatures for vaccine maintenance globally is between 2°C and 8°C (31,32). It is generally recommended that vaccines should be stored in their original packages and, for lyophilized vaccines, to only be reconstituted when needed. In this study, it was found that some agro-shops were reconstituting vaccines and drawing them into syringes for sale to customers. Reconstituted vaccines are more sensitive than non-reconstituted vaccines and are at a higher risk of bacterial contamination and overgrowth if the syringes are left or not administered for prolonged periods. The practice of selling reconstituted vaccines compromises the quality and potency of vaccines. Furthermore, the sale of reconstituted vaccines denies customers their right to product information and increases the risk of malpractices due to lack of product identity.

Thermostable I-2 ND vaccines were developed to reduce dependence on cold chains. The strain I-2 of ND vaccine has many advantages that include thermostability, easy administration by various routes, such as drinking, eye drop, and mixing with food, and providing good protection against the disease (33). The vaccines have been recommended for use in village chicken as they can remain stable and potent outside the recommended cold chain temperatures for a considerable time. However, these vaccines were not widely available as the results of this study show. The most common type of vaccines found in the agro-shops in this study were the thermolabile Lasota types despite the fact they were relatively more expensive and require more stringent cold chain conditions.

Guidelines and best practices in the handling of veterinary vaccines, including the Newcastle disease vaccine, have been published and are readily available (30). However, some of these may have limited application in resource-poor settings, as suggested by the results of this study. For most agro-shops in rural areas, providing quality and potent poultry vaccines to customers, the majority of whom are smallholder farmers, in an environment that is characterized by frequent power outages and escalating costs of electricity is no easy task. As the results of this study show, most agro-shops had cold chain equipment for the storage of vaccines. While this is essential, it does not automatically result in optimal vaccine storage conditions as improperly maintained or outdated refrigeration equipment, poor compliance with cold-chain procedures, and inadequate temperature monitoring could affect cold storage conditions (18).

## Conclusion

The results of the present study have revealed several practices in the last mile, transportation, storage and sale of vaccines by agro-shops. Faced with long-standing challenges in the maintenance of cold chains, the agro-shops devised several practices that may be affecting the quality and potency of vaccines in one way or another. These practices include restricting the sale of vaccines to late hours of the afternoon when the ambient temperature is lower and periodically switching off the fridge and restricting the hours of sale of vaccines to minimize storage costs. However, it remains unclear how such practices affect vaccine quality and potency. Therefore, further research could provide more information on the effects of such practices on vaccine potency.

Thermostable vaccine formulations can leave cold-chain conditions for a comparatively longer time while retaining potency. Thermostability of vaccines is a desirable trait in resource-poor settings, particularly where maintenance of cold chain is difficult. However, as the results of this study showed, thermo-stable vaccines were not widely available in the agro-shops. It has been suggested that where there is an existing, effective, low-cost vaccine in routine use, it is unlikely that manufacturers would change their formulations or manufacturing processes for an established low-profit-margin vaccine (34). However, further research could provide more information for the apparent preference of lyophilized vaccines which require more stringent cold chain conditions.

Agro-shops are a critical link in the delivery of Newcastle disease and other poultry vaccines to smallholder farmers in rural areas. Proper transport, storage, and handling of vaccines are issues that are frequently overlooked when creating or implementing vaccine protocols and vaccination programs. Between the time a vaccine leaves the manufacturer’s plant and the time it is injected into an animal, there are many stages for inadvertent contamination or inactivation. By being aware of these potential “weak points” in a vaccine protocol, agro-shops can help ensure that vaccines are not rendered ineffective because of improper handling. Good practices to maintain proper vaccine storage and handling can ensure that the full benefit of immunization is realized.

## Acknowledgment

The authors would like to acknowledge all the agro-shops and enumerators that participated in the study and the County Government of Bungoma for their help and cooperation.

